# Reconstructing Genetic History of Siberian and Northeastern European Populations

**DOI:** 10.1101/029421

**Authors:** Emily HM Wong, Andrey Khrunin, Larissa Nichols, Dmitry Pushkarev, Denis Khokhrin, Dmitry Verbenko, Oleg Evgrafov, James Knowles, John Novembre, Svetlana Limborska, Anton Valouev

## Abstract

Siberia and Western Russia are home to over 40 culturally and linguistically diverse indigenous ethnic groups. Yet, genetic variation of peoples from this region is largely uncharacterized. We present whole-genome sequencing data from 28 individuals belonging to 14 distinct indigenous populations from that region. We combine these datasets with additional 32 modern-day and 15 ancient human genomes to build and compare autosomal, Y-DNA and mtDNA trees. Our results provide new links between modern and ancient inhabitants of Eurasia. Siberians share 38% of ancestry with descendants of the 45,000-year-old Ust’-Ishim people, who were previously believed to have no modern-day descendants. Western Siberians trace 57% of their ancestry to the Ancient North Eurasians, represented by the 24,000-year-old Siberian Mal’ta boy. In addition, Siberians admixtures are present in lineages represented by Eastern European hunter-gatherers from Samara, Karelia, Hungary and Sweden (from 8,000-6,600 years ago), as well as Yamnaya culture people (5,300-4,700 years ago) and modern-day northeastern Europeans. These results provide new evidence of ancient gene flow from Siberia into Europe.

## INTRODUCTION

Understanding the population history of Siberian and the Trans-Uralic region (the territory to the West and East of the Ural Mountains) is of great historical interest and would shed light on the origins of modern-day Eurasians and populations of the New World. Large-scale population genetic mapping efforts such as 1000 Genomes^1^ and HapMap^2^ have not analyzed populations from these regions despite their prominent role in peopling of Eurasia^3,4^ and America^5^

Siberia has been inhabited by hominids for hundreds of thousands of years, with some of the known archeological sites being older than 260,000 years^6^. Neanderthals inhabited Europe and Siberia until approximately 40,000 years ago^7^, when anatomically modern humans expanded to that region 60,000–40,000 years ago^8^. A high-coverage genome of a Siberian hominine individual, known as Denisovan, was recently analyzed and compared to the related Neanderthal genome^9^. The age of Denisovan was estimated to be 74,000-82,000 years^10^. Traces of habitation of anatomically modern humans in Siberia date to at least 45,000 years ago, based on a bone discovered recently near Ust’-Ishim settlement in Western Siberia^11^. Ust’-Ishim’s genome carried genomic segments attributed to admixtures with Neanderthal, estimated to have occurred 52,000-58,000 years ago based on collagen dating and genetic methods. Other ancient human sites in Siberia yielded ancient DNA from Ancient North Eurasians, or ANE^4^, including the Upper Paleolithic 24,000-year-old Siberian Mal’ta boy MA-1^5^ and the 17,000-year-old Siberian AG-2^5^. Genetic analysis of these samples showed that 42% of the Native American genetic makeup could be traced to the ANE people related to Mal’ta boy. Yet our understanding of genetic origins of present-day indigenous Siberians and their relationships with ancient populations is still far from being complete.

### Western Siberians and Northeastern Europeans

Modern Siberian and Eastern European populations harbor enormous genetic and cultural diversity. Indigenous people of the Trans-Uralic region speak languages broadly categorized as Uralic, which are further subdivided into Ugric, Finno-Permic and Samoyedic groups (Supp. Fig. 1). Ugric languages are spoken by two small Western Siberian groups of Khanty and Mansi as well as 13 million Hungarians in Central Europe. Ugric languages are distantly related to Permic and Finno-Volgaic language groups, native to several populations in Northeastern Europe, including Komi, Karelians, Veps, Saami and Finns. The language of northwestern Siberian Nenets people belongs to the Samoyedic language group, which also includes languages spoken by small western Siberian populations of Enets and Nganasan people.

### Eastern Siberians

Languages of Central and Eastern Siberian populations are broadly categorized as Altaic, and are further subdivided into Mongolic, Tungusic and Turkic groups. The Buryat and Kalmyk people are members of the Mongolic language group. The Even and Evenki Tungusic languages are spoken over vast but sparsely populated regions of Central and Eastern Siberia. Tungusic languages used to be much more widespread and were previously spoken by the Manchu people, who founded the Qing dynasty but later adopted Mandarin Chinese. The Altayan and Yakut populations are the most numerous Siberian groups, who speak Siberian Turkic languages.

### Whole-genome sequencing

We performed deep sequencing of 28 individuals, representing 14 ethnic groups from Siberia and Eastern Europe (Table 1, Fig. 1a). To maximize the quality of the population data, we obtained DNA samples that are informative for deep population history of individuals from geographical locations traditionally occupied by the corresponding indigenous populations and without self-reported mixed ancestry for at least three generations. Our samples cover major populations from Siberia and Eastern Europe, representing speakers of primary linguistic groups from these regions.

**Table 1.** List of genomes from Siberian and Eastern European populations

| **Ethnic group** | **Samples** | **Census size (M = 10<sup>6</sup>)** | **Language group** | **Geographic distribution** | **Sequence coverage** | **Non-dbSNP SNPs (in thousands)** |
| --- | --- | --- | --- | --- | --- | --- |
| Komi Izhma | 2 | 225,000 | Finno-Permic | Eastern Europe | 37x, 46x | 74, 85 |
| Komi Ob'yachevo | 2 | 225,000 | Finno-Permic | Eastern Europe | 31x, 37x | 82, 85 |
| Veps | 2 | 6,400 | Finno-Permic | N.-E. Europe | 39x, 42x | 75, 76 |
| Karelians | 1 | 89,000 | Finno-Permic | N.-E. Europe | 36x | 209 |
| Russians (Mezen) | 2 | 111 M | Slavic | N.-E. Europe | 33x, 38x | 69, 71 |
| Russians (Ustyuzhna) | 2 | 111 M | Slavic | Eastern Europe | 40x, 50x | 246, 424 |
| Russians (Andreapol) | 2 | 111 M | Slavic | Eastern Europe | 36x, 46x | 238, 412 |
| Belarusians | 1 | 8 M | Slavic | Eastern Europe | 30x | 204 |
| Mansi | 3 | 12,500 | Ugric | Western Siberia | 44x, 45x, 47x | 222, 229, 244 |
| Khanty | 4 | 31,500 | Ugric | Western Siberia | 31x, 39x, 43x, 44x | 92, 85, 313, 319 |
| Nenets | 1 | 45,000 | Samoyedic | N.-W. Siberia | 43x | 90 |
| Evens | 1 | 22,000 | Tungusic | Central Siberia | 34x | 151 |
| Evenki | 1 | 80,000 | Tungusic | Eastern Siberia | 33x | 157 |
| Yakuts | 1 | 480,000 | Turkic | Eastern Siberia | 33x | 179 |
| Altayans | 1 | 70,800 | Turkic | Central Siberia | 35x | 166 |
| Buryats | 1 | 600,000 | Mongolic | Central Siberia | 31x | 152 |
| Kalmyks | 1 | 183,000 | Mongolic | S.-E. Europe | 33x | 152 |

**Figure 1.**
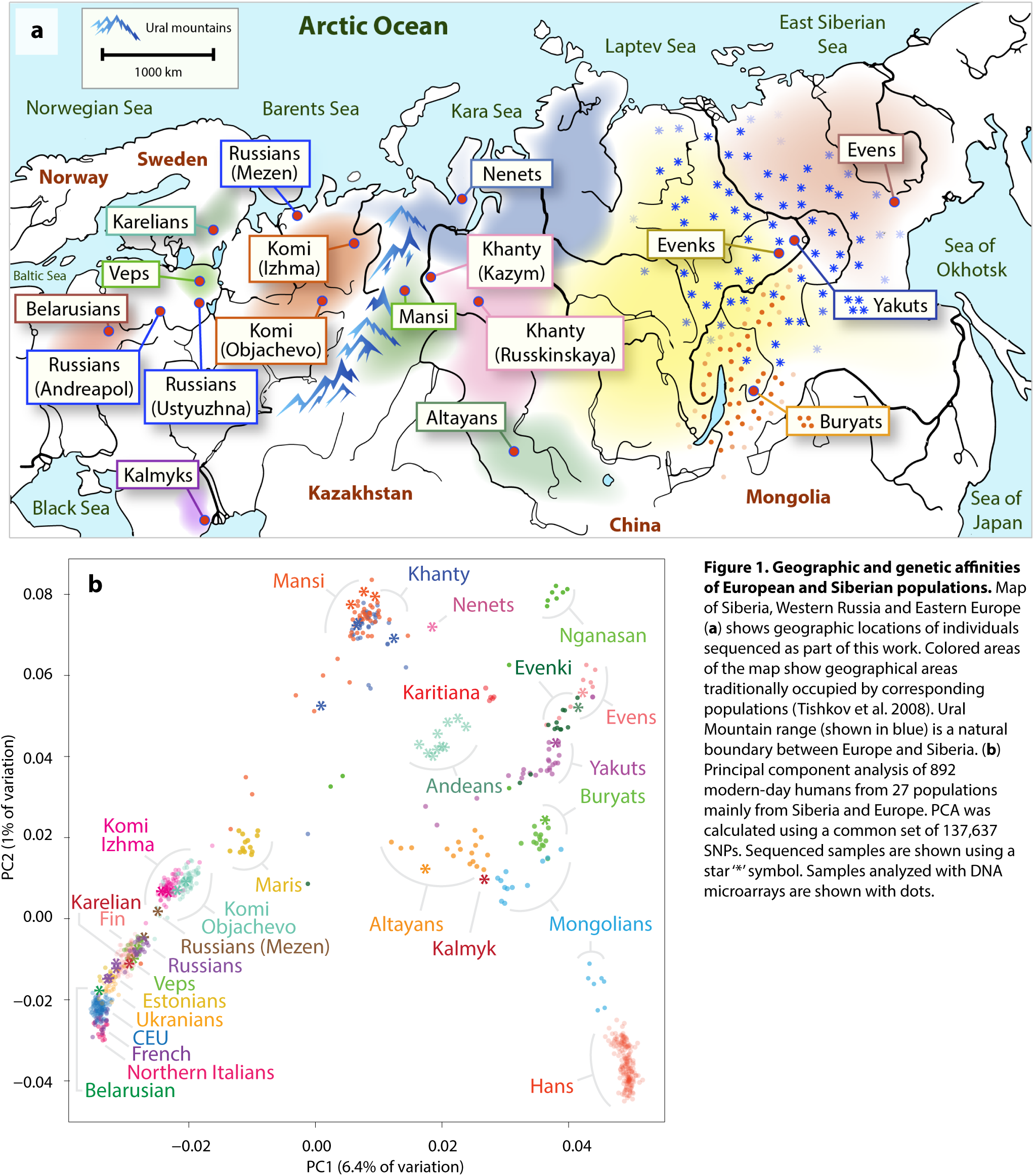
Geographic and genetic affinities of European and Siberian populations. Map of Siberia, Western Russia and Eastern Europe (**a**) shows geographic locations of individuals sequenced as part of this work. Colored areas of the map show geographical areas traditionally occupied by corresponding populations (Tishkov et al. 2008). Ural Mountain range (shown in blue) is a natural boundary between Europe and Siberia. (**b**) Principal component analysis of 892 modern-day humans from 27 populations mainly from Siberia and Europe. PCA was calculated using a common set of 137,637 SNPs. Sequenced samples are shown using a star ‘*’ symbol. Samples analyzed with DNA microarrays are shown with dots.

DNA samples were collected with informed consent under the IRB-equivalent approval from the Institute of Molecular Genetics, Russian Academy of Sciences. Samples were further de-identified and analyzed under IRB approval (HS-13-00594) from the University of Southern California. All samples were sequenced on the Illumina platform to high-depth of at least 30X (based on uniquely mapped reads) with the average sequence coverage depth of 38X. All genomes were analyzed using the same set of computational tools (Novoalign and GATK^12^) and filtering criteria (Supplementary Information) to obtain high-confidence SNP calls. In addition, we genotyped samples from two Siberian populations, Mansi and Khanty (45 and 37 individuals respectively), using high-density SNP microarrays (Supp. Information).

### Population relationships

First, we sought to obtain a broad overview of the population genetic relationships using PLINK^13^ and EIGENSOFT^14,15^ for Principal Component Analysis (PCA). We compared 892 present-day humans^1,5,16-20^ from 27 Asian, European, Siberian and Native American populations using a common set of 137,639 autosomal SNP loci (Fig. 1b; Supp. Fig. 2; Sup. Table 1). This analysis included the 82 new samples from Mansi (N=45) and Khanty (N=37) groups that were genotyped across 628,515 SNPs using Illumina microarrays. Major differentiation of populations is captured by the first principal component, which mimics West-to-East distribution of people across East Asia and Siberia and explains 6.4% of genetic variation. The second principal component reflects the spread of populations along the North-to-South latitudinal cline, especially among Siberians, capturing 1% of genetic variation. The PCA plot reveals a significant degree of genetic differentiation between Siberian groups, while European populations show only modest levels of population-specific variation.

### Autosomal phylogeny

To examine the genetic relationships of Siberian and Eastern European populations relative to other populations in the world, we integrated publicly available raw sequencing data from 32 high-coverage modern genomes representing 18 populations^9,17,21-23^ (Supp. Table 2) as well as variant calls from the two hominin genomes, the Neanderthal^9^ and Denisova individuals^10^ in our analysis. We sought to minimize potential biases stemming from the use of different sequencing platforms, different read mapping tools, different SNP calling tools and downstream variant filters. Therefore, all genomes (except for the Denisova and Neanderthal samples) were re-analyzed starting from raw sequencing reads using the same set of tools, parameters and filtering criteria as the other genomes in this study (Supp. Information). After SNP calling and filtering, we constructed a full autosomal TreeMix^24^ admixture graph (Supp. Fig. 3-4) using a set of 25,589,077 genome-wide high-quality autosomal SNPs. TreeMix models demographic scenarios in the form of a bifurcating tree, allowing for admixture events between individuals and populations to provide insights into hidden demographic events of the past.

Consistent with the model of out-of-Africa human migration into other parts of the world, the tree model places African populations as outgroups relative to Eurasian and Native American people. Furthermore, non-African populations are clustered into two primary groups: one group that includes South Indians and all European populations (except for Kalmyks, who migrated to the lower Volga region in Eastern Europe from Dzungaria, a region in northwestern China, in 17th century^25^), and one that encompasses Papuans, Native Australians, all Asian and Siberian people. The inferred admixture events further suggest that significant gene flow exists between South Indians and Malaysians as well as South Indians and Native Australians, tracing 8% (95% CI: 5-10%) of the South Indian ancestry to the population ancestral to Malaysians and 19% (95% CI: 17-20%) to Native Australians. Furthermore, Han, Dai, Sherpa and Malaysians, the East Asian populations, have significant admixture with peoples related to Australian Aboriginals accounting for 19% of their ancestry. Consistent with previous reports^5,26^, we also observed gene flow from a population related to Denisovans and Neanderthals into the common ancestor of Papuans and Australian Aboriginals accounting for 18% (95% CI: 17-20%) of their ancestry. This estimate is higher than the previously reported 3-8% admixture, which we also reproduced using a smaller set of only eight modern individuals (Supp. Fig. 6). The higher estimate is also robust to using other methods such as F4-ratio test14, which indicates that 17-21% of Australian Aboriginal ancestry and 26-30% of Papuan ancestry is hominine-related (Supp. Fig. 24). The higher percentage of hominine-related admixture among Oceanians reported here (Supp. Fig. 3) is likely explained by a more accurate phylogenetic TreeMix model, since it incorporates nearly an order of magnitude more individuals compared to previous studies, including Siberian populations, which were not previously used for this inference.

To reveal weaker admixture events within Siberian and European groups, we excluded the admixed Papuan, Native Australians, South Indians and Malaysian individuals and recalculated the tree model (Fig. 2). In this phylogenetic inference, the Western Siberian populations of Khanty, Mansi and Nenets appear as an early diverging group related to other European populations. The tree model suggests that 43% (95% CI: 38-47%) of the Western Siberian ancestry can be attributed to an admixture with a group related to modern-day Evenki people. Furthermore, Nenets share 38% (95% CI: 31-46%) of their ancestry with a group related to Even people. Consistent with this prediction, we observed particularly high affinity between Mansi and Evenki as well as between Nenets and Even people based on the D-statistic^27,28^ (Supp. Fig. 7a-b, 8a-b). TreeMix inferred admixture between common ancestors of Mansi, Khanty and Nenets and Andean Highlanders that accounts for 6% (95% CI: 4-8%) of the Western Siberian ancestry. This admixture can be explained by a lineage of Ancient North Eurasians ancestral to both Western Siberians and Native Americans and is discussed later.

**Figure 2.**
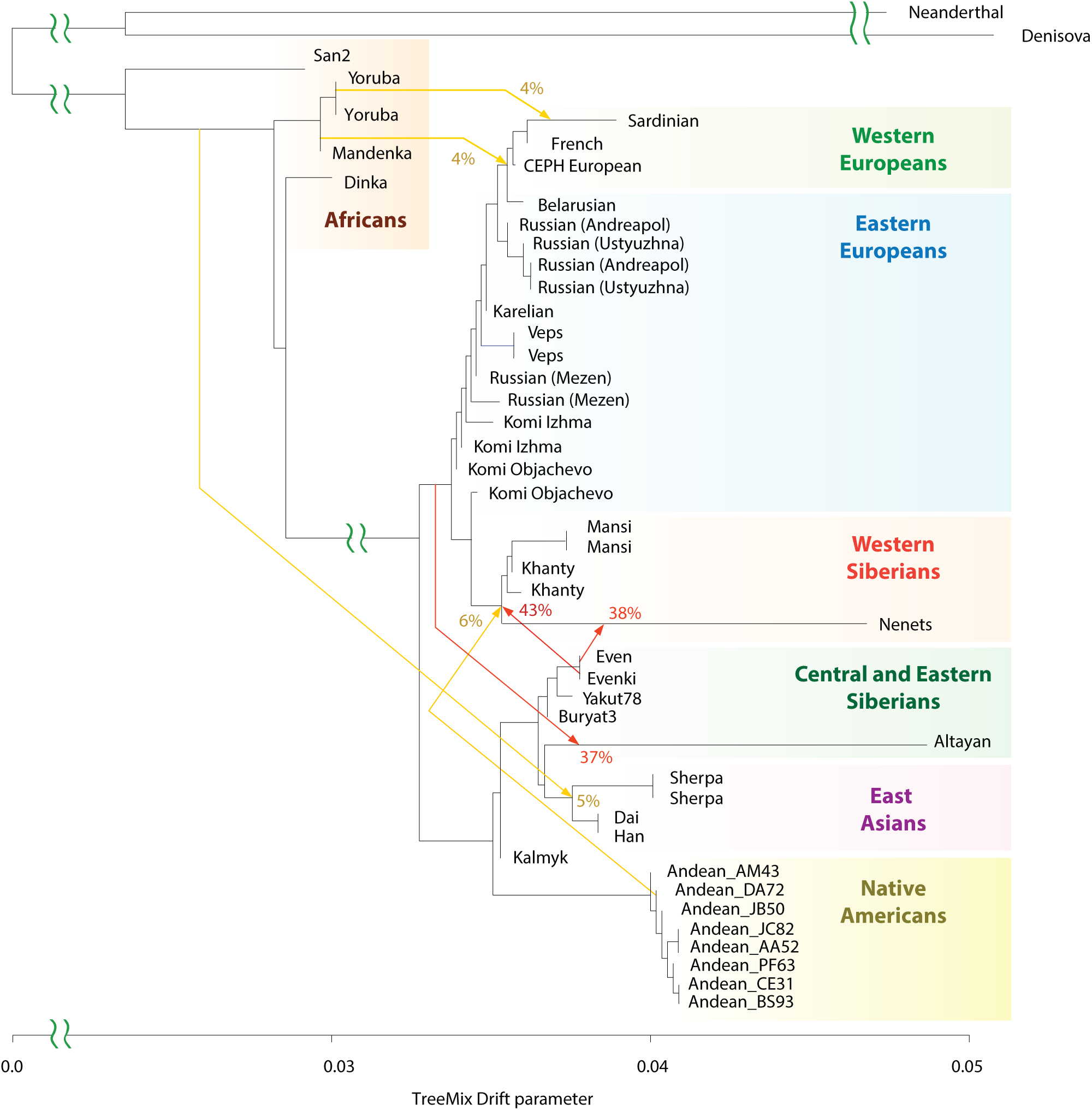
Genetic relationships of modern-day Siberian, Asian, Native American, European and African populations. Autosomal TreeMix admixture graph based on whole-genome sequencing data from 44 individuals primarily from Africa, Europe, Siberia and America. The X-axis represents genetic drift, which is proportional to the effective population size N_e_. Admixture events are shown with arrows, with admixture intensities indicated by percentages. Residuals are shown in Supp. Fig. 5.

Phylogenetic trees provide simplified demographic models. The lengths of branches are expressed in the units of drift parameter, and can be affected by population bottlenecks, further complicating the inference of population divergence time. Therefore, we used an alternative MSMC method^29^ (Supp. Information) that calculates separation times between pairs of populations (Fig. 3a). As a proxy for separation time of two populations, we utilized the time at which the relevant cross-coalescence rate became 0.5 (Fig. 3b).

**Figure 3.**
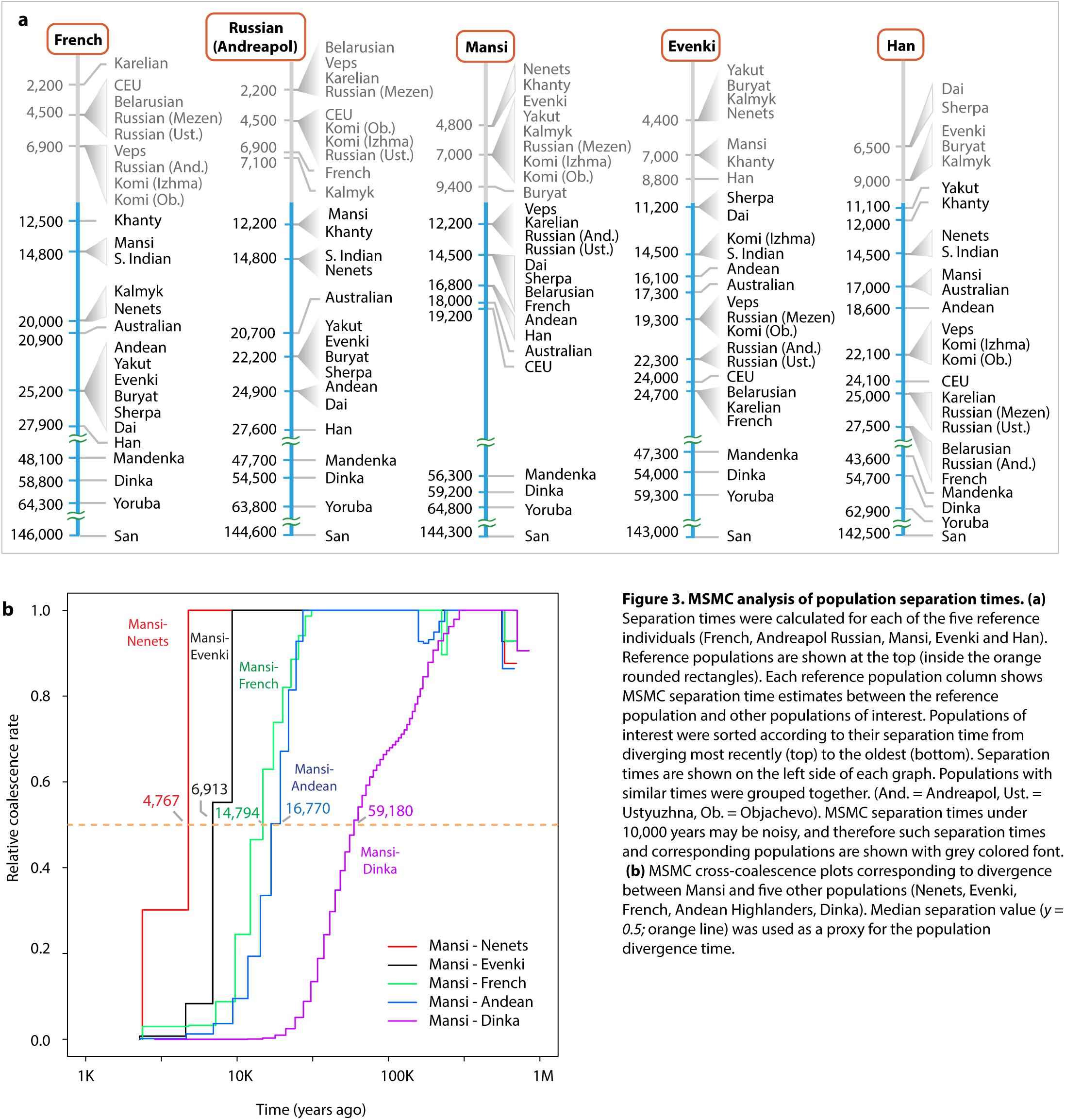
MSMC analysis of population separation times. (**a**) Separation times were calculated for each of the five reference individuals (French, Andreapol Russian, Mansi, Evenki and Han). Reference populations are shown at the top (inside the orange rounded rectangles). Each reference population column shows MSMC separation time estimates between the reference population and other populations of interest. Populations of interest were sorted according to their separation time from diverging most recently (top) to the oldest (bottom). Separation times are shown on the left side of each graph. Populations with similar times were grouped together. (And. = Andreapol, Ust. = Ustyuzhna, Ob. = Objachevo). MSMC separation times under 10,000 years may be noisy, and therefore such separation times and corresponding populations are shown with grey colored font. **(b)** MSMC cross-coalescence plots corresponding to divergence between Mansi and five other populations (Nenets, Evenki, French, Andean Highlanders, Dinka). Median separation value (*y = 0.5;* orange line) was used as a proxy for the population divergence time.

The MSMC results suggested that Mansi, Khanty and Nenets separated relatively recently (4.8 thousand years ago or kya) after experiencing admixture with Evenki group (Fig. 2). The separation between Mansi and Evenki occurred approximately 6.8 kya, indicating that Eastern Siberian admixture into Western Siberian groups occurred within the range of 4.8-6.8 kya. The 4 haplotype MSMC estimates, based on one individual from each population, should be interpreted with caution, as MSMC results may be noisy for separation times below 10,000 years^29^. However, the relative order of populations is still informative for interpreting their relative genetic affinities. The more accurate 8 haplotype MSMC analysis, provides a similar estimate of 5-9.9 kya for the time of the Eastern Siberian admixture into Western Siberian populations (Supp. Fig. 9).

The phylogenetic tree model placed all modern-day Europeans (except for Kalmyk) along a single branch, with the divergence pattern mimicking the East-to-West distribution of populations within Europe. The separation times between French and Eastern European populations fell in the range of 2.3 kya (French-Karelian) to 7 kya (French-Komi Objachevo), confirming the observation from the PCA analysis (Fig. 1b) that Western and Eastern European populations were fairly close genetically.

Siberian, Native American and Asian populations, as well as Kalmyk, form a second major clade within the tree. Notably, Central and Eastern Siberian populations, such as Buryat, Yakut, Even, Evenki and Altayan form a distinct sub-lineage, after diverging from Han, Dai and Sherpa, the East Asian populations. The separate clade of Eastern Siberians is also supported by a larger tree, which also includes Malaysians, who group with Han and Dai (Supp. Fig. 3). We estimated the separation time between Eastern Siberians and East Asian populations to be within the range between 8.8 kya (Evenki-Han) and 11.2 kya (Evenki-Sherpa). The relationship of Kalmyk and Altayan indivduals with the respect to other Siberians and East Asians was not apparent from their positions on the autosomal tree. Based on the D-statistic^27,28^, both Kalmyk and Altayan are moderately closer to other Siberians such as Even and Evenki (D-statistics = 0.13-0.146; Supp. Fig. 7c-d, 8c-d) than to East Asians (D statistics = 0.11-0.125). However, unlike Kalmyk, Altayans likely trace approximately 37% (95% CI: 31-43%) of their ancestry to another unknown population, which the model predicted to be related to modern Europeans (Fig. 2).

### Y-DNA and mtDNA analyses

To obtain an independent set of population divergence time estimates, we calculated divergence times of Y-chromosome haplotypes using SNP calls from the genomes of 54 modern-day individuals representing 30 world populations. Both Y-DNA and mtDNA accumulate derived alleles (or mutations) within a single haplotype at known mutation rates, thus allowing inference of haplotype divergence times. We utilized SNPs within unique non-homologous regions of Chromosome Y totaling 10.4 Mb^30^ and a mutation rate of 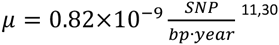 to estimate the haplotype divergence times (Methods; Fig. 4; Supp. Fig. 10-11). To investigate divergence of mtDNA haplotypes, we used the data from 70 individuals and 34 world populations and a mutation rate of 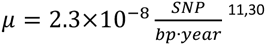. In addition to modern-day genomes, we also placed eight ancient individuals^4,5,31-35^ (Supp. Table 3) on the calculated haplogroup trees according to the sharing pattern of high-confidence derived alleles inferred to arise along each branch.

**Figure 4.**
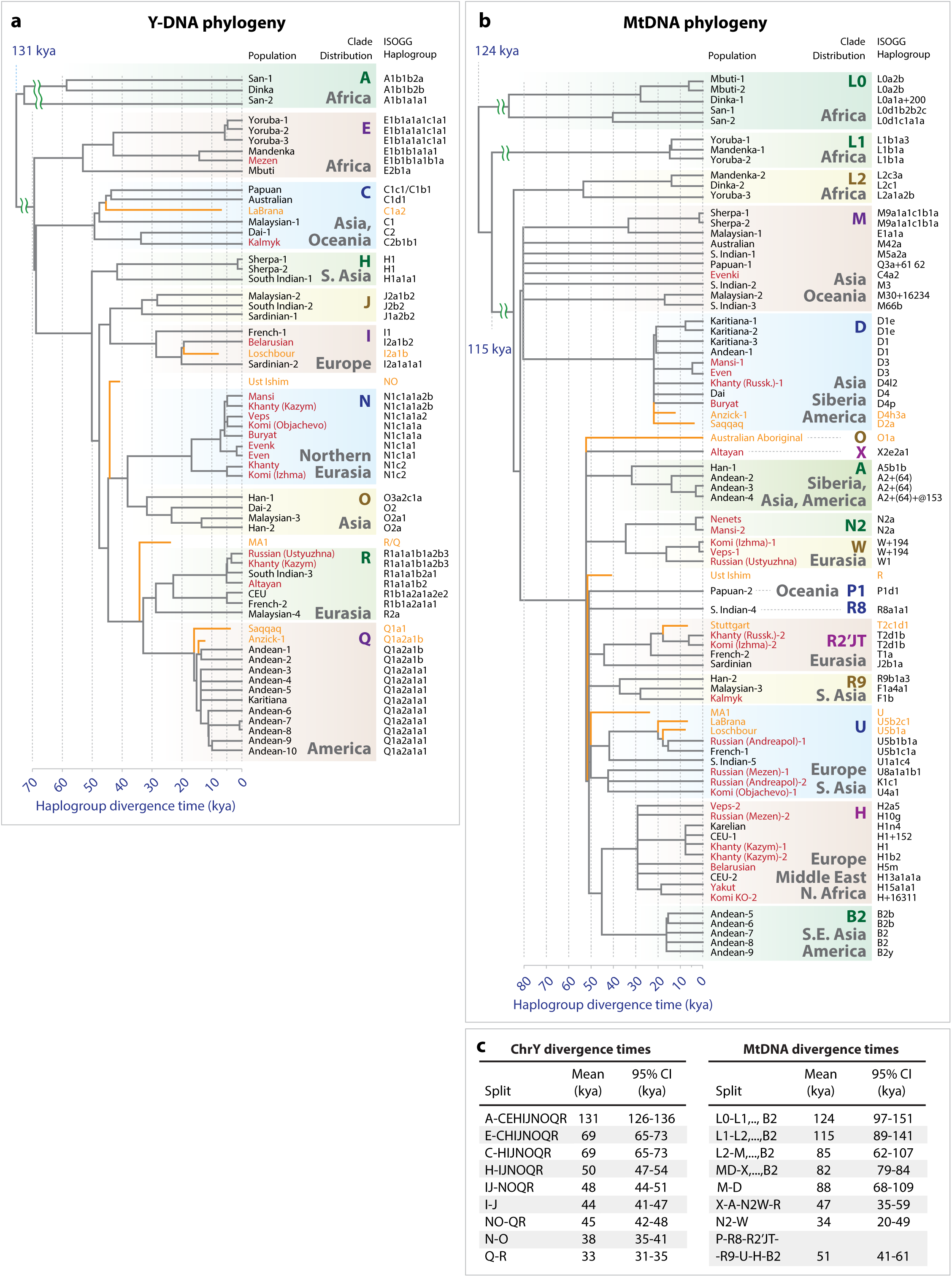
Y-DNA and MtDNA phylogeny and haplogroup divergence times. **(a)** Y-DNA tree topology was calculated using maximum likelihood approach implemented in MEGA tool (Tamura et al. 2011) using 12,394 polymorphic sites inferred from SNP calls. The X-axis represents divergence time of haplogroups and is expressed in units of thousands of years ago (1 kya = 1,000 years ago). The tree was constructed using the data from 54 modern-day humans from 30 populations. Split times were calculated according to the number of derived alleles assigned to each branch (Methods) and splits were placed along the X-axis to reflect haplotype divergence time. Terminal branches end at X=0 for modern-day individuals. Seven ancient individuals were placed on the haplotype tree according to the SNP-sharing patterns with high-confidence derived allele SNVs assigned to each branch. Branches of ancient individuals (orange lines) terminate at times when each ancient individual lived. Haplogroups were assigned based on ISOGG Y-DNA SNP index and shown along the right side of the tree (ISOGG, 2015). Individuals sequenced as part of this work are shown in red. **(b)** MtDNA phylogeny was constructed using MEGA tool in a similar fashion to Y-DNA tree. The tree includes data from 70 individuals and 34 populations as well as 8 ancient genomes. **(c)** Divergence times of haplogroups from Y-DNA phylogeny (left) and MtDNA phylogeny (right). Diverging clades are shown on the left-hand side (“Split” column). Confidence interval were calculated using Poisson model.

The earliest diverging Y-chromosome lineage in our dataset is haplogroup A, carried by two African San and a Dinka individual. The age of this lineage is 131 kya (Fig. 4; 95% CI: 126-136 kya), which is consistent with a recent estimate of 120-156 kya^30^ and 115-157 kya^11^. Haplogroup E is closest to Eurasian clades diverging 69 kya (95% CI: 65-73 kya), providing an upper bound time estimate for the out-of-Africa migration.

Within the mtDNA tree, the earliest diverging haplogroup is L0, which has an age of 124 kya (95% CI: 97-151 kya). This matches previous estimate of 124 kya^30^ (95% CI: 99-148 kya). The mtDNA data shows three initially diverging clades L0, L1 and L2, primarily found within African populations, with most recently diverging L2 clade having age of 85 kya (95% CI: 62-107). Therefore estimates of out-of-Africa migration time based on Y-DNA and mtDNA phylogenies are slightly higher than MSMC separation time estimates of 44-56 kya (Mandenka) or 55-59 kya (Dinka). The MSMC estimate only provides an average separation between two populations, without specifically accounting for admixtures. Therefore, the MSMC separation time estimate is expected to be lower than the actual divergence time of two populations due subsequent admixtures and gene flow events following their initial divergence.

The divergence of haplotypes carried by Native Americans Karitiana (Brazil) and Andean Highlanders (Peru) is consistent with initial colonization of America 15,000 years ago^36^. The Y-chromosome data indicates that the divergence of Native American haplogroup Q from the Eurasian haplogroup R occurred approximately 33 kya (95% CI: 30-36 kya), as expected, predating the arrival of Paleo-Indians into America 15,000 years ago. The expansion of haplogroup subclades originating within Native American haplogroups is expected to postdate the time of the expansion into the New World. In agreement with this prediction, we estimated the expansion time of the Q clade lineage to be 15.1 kya (95% CI: 13.3-17.0 kya). We made similar observations based on mtDNA data. Haplogroups from three mtDNA clades D1, A2 and B2 are common among Native Americans. We estimated that the divergence of Native American and Eurasian haplogroups occurred 22 kya (95% CI: 15-29; clade D), 32 kya (95% CI: 18-45; clade A) and 45 kya (95% CI: 29-62; B2-H clade split). The expansions of Native American mitochondrial clades occurred 13.9 kya (95% CI: 4.8-23; clade A2), 14.7 kya (95% CI: 8-21; clade B2) and 24.3 kya (95% CI: 14-34; clade D1).

Major Y-chromosome clades common among Europeans include I, J, R and N. Among those, haplogroup I is the oldest surviving clade with mainly European distribution, diverging from its sister clade J 44 kya (95% CI: 41-47 kya) and expanding within Europe at least 29 kya. Diverging from IJ clade 48 kya (95% CI: 44-51) is clade K2 (formely K(xLT)), which is ancestral to clades NO (or K2a) and QR (or K2b). Ancient Siberian Ust’-Ishim^11^ is thought to belong to K2 Y-DNA clade. We observed that all 16 K2 – specific SNPs were present in Ust’-Ishim. However, Ust’-Ishim also shares one out of 47 derived alleles specific to NO branch (hg19 coordinate ChrY 7690182; Supp. Fig. 12). Therefore, Ust’-Ishim’ s haplogroup most likely belongs to NO clade that diverged shortly after NO-QR split (45 kya; 95% CI: 42-48 kya), but before N-O split (38 kya; 95% CI: 35-41). We also calculated that Ust’-Ishim’s Y-DNA haplogroup had 352 and 335 fewer private SNPs relative to what was expected for a modern-day individual (based on QR and NO clade individuals respectively). This provides two independent estimates of 41,100 years (95% CI: 37,100-45,000 years) and 39,200 years (95% CI: 35,600-42,900) for Ust’-Ishim’s age.

Within Europe, clade R further diverges into two subclades R1a1 and R1b1 23 kya. Clade R1b1 is primarily found among Western Europeans. Its sister clade R1a1 is common across Eastern Europe, but one of its subclades R1a1a1b2 is also common within Altay region as well as in Northern India and Pakistan^37^. Our Y-chromosome haplogroup tree shows that South Indian and Altayan individuals share a common R1a1a1-haplogroup ancestor along the male lineage approximately 5.1 kya, suggesting ancient migration routes connecting Eastern Europe, Altay region and India.

The Y-chromosome haplogroup N is spread across Siberia and Eastern Europe and reaches its maximum frequency among Siberian populations such as Yakut (90%)^38^, Nenets (97%)^39^ and Nganasan (92%)^39^. Among Eastern Europeans, it is mainly found in certain Russian groups (71%)^40,41^ and Komi (70-75%)^42^, and is nearly absent in Western Europeans. This suggests that haplogroup N reached Europe only relatively recently and had not spread yet to Western Europe. The split between clades N and O occurred approximately 38.2 kya (95% CI: 35-41 kya), followed by formation of two subclades N1c1 and N1c2 clades 16.8 kya. N1c1, the more common clade, is widely spread in both Siberia and Eastern Europe, where it reaches high frequency among Finns (58%)^43^ and Saami (47%)^39^ populations. This cluster expanded 5.3-7.1 kya, and spread within Siberians (Even, Evenki, Mansi, Khanty) and Eastern Europeans (Veps and Komi). This suggests the existence of gene flow from Siberian populations into Eastern and Northern Europe, which occurred approximately 5.3-7.1 kya.

We investigated this hypothesis further, by analyzing gene flows occurring within approximately the last 7,000 years in northeastern European populations of Mezen Russians, Veps, Karelians, and Komi (Fig. 5a-b, Supp. Fig. 7f,j,k, Supp. Fig. 13a-b). The northeastern Europeans show statistically significant admixture signals with Siberian groups such as Yakut, Nenets, Even, and Khanty. These admixtures are much weaker among more southeastern Europeans Ustyuzhna and Andreapol Russians as well as Belarusian individual (Supp. Fig. 7g-I, Supp. Fig. 13g-i), suggesting that Siberian admixtures are particularly strong among northeastern Europeans.

**Figure 5.**
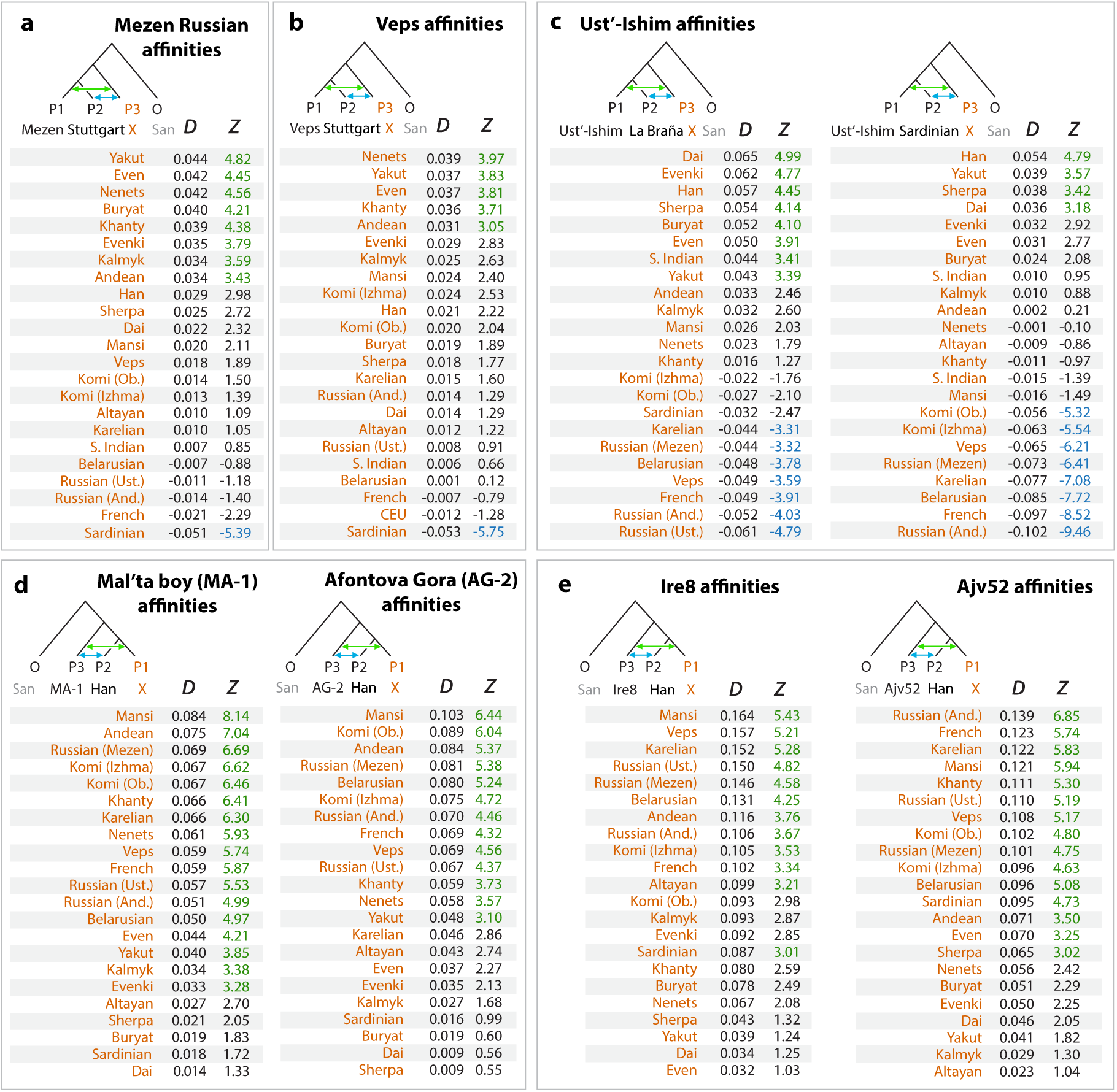
Analysis of admixtures using D-statistics. Genetic affinities were determined using *D*-statistics test *D(P_1_, P_2_, P_3_, O*) from the ADMIXTOOLS package (Patterson et al. 2012) and shown under *D* column. Statistical significance of admixtures was expressed in terms of z-score units and shown under *Z* column. Positive D-statistics values indicate possible admixture between *P_1_* and *P_3_*_′_ while negative - between *P_2_* and *P_3_*_′_ Green font was used to highlight significant admixtures between *P_1_* and *P_3_* (*Z>3*). Blue font was used to highlight significant admixture between *P_2_* and *P_3_* (*Z<-3*). African San was used as an outgroup in all tests. (And. = Andreapol, Ust. = Ustyuzhna, Ob. = Objachevo). D-statistics results using different outgroups (Chimp, Mandenka, Mbuti) are shown in Supp. Fig. 8–10.

### Comparison with ancient genomes

Examining genetic affinities relative to ancient genomes uncovers ancient demographic events and reveals genetic links between present-day individuals and ancient ancestral populations. We considered 15 ancient genomes (Supp. Table 3), including a 45,000-year-old Siberian Ust’-Ishim^11^, a 24,000-year-old Siberian Mal’ta boy (MA-1) and a 17,000-year-old Siberian individual (AG-2) from the Krasnoyarsk region^5^.

The TreeMix model placed the Ust’-Ishim individual as an outgroup relative to the split between European and Asian clades (Fig. 6, Supp. Fig. 14). Furthermore, the common ancestor of Siberians and East Asians traces 38% (95% CI: 28-48%) of the ancestry to Ust’-Ishim’ s lineage. In agreement with this, the D-statistic (Fig. 5c, Supp. Fig. 13c, Supp. Fig. 15) showed that East Asians and Eastern Siberians had higher genetic affinity with Ust’-Ishim than modern Europeans.

**Figure 6.**
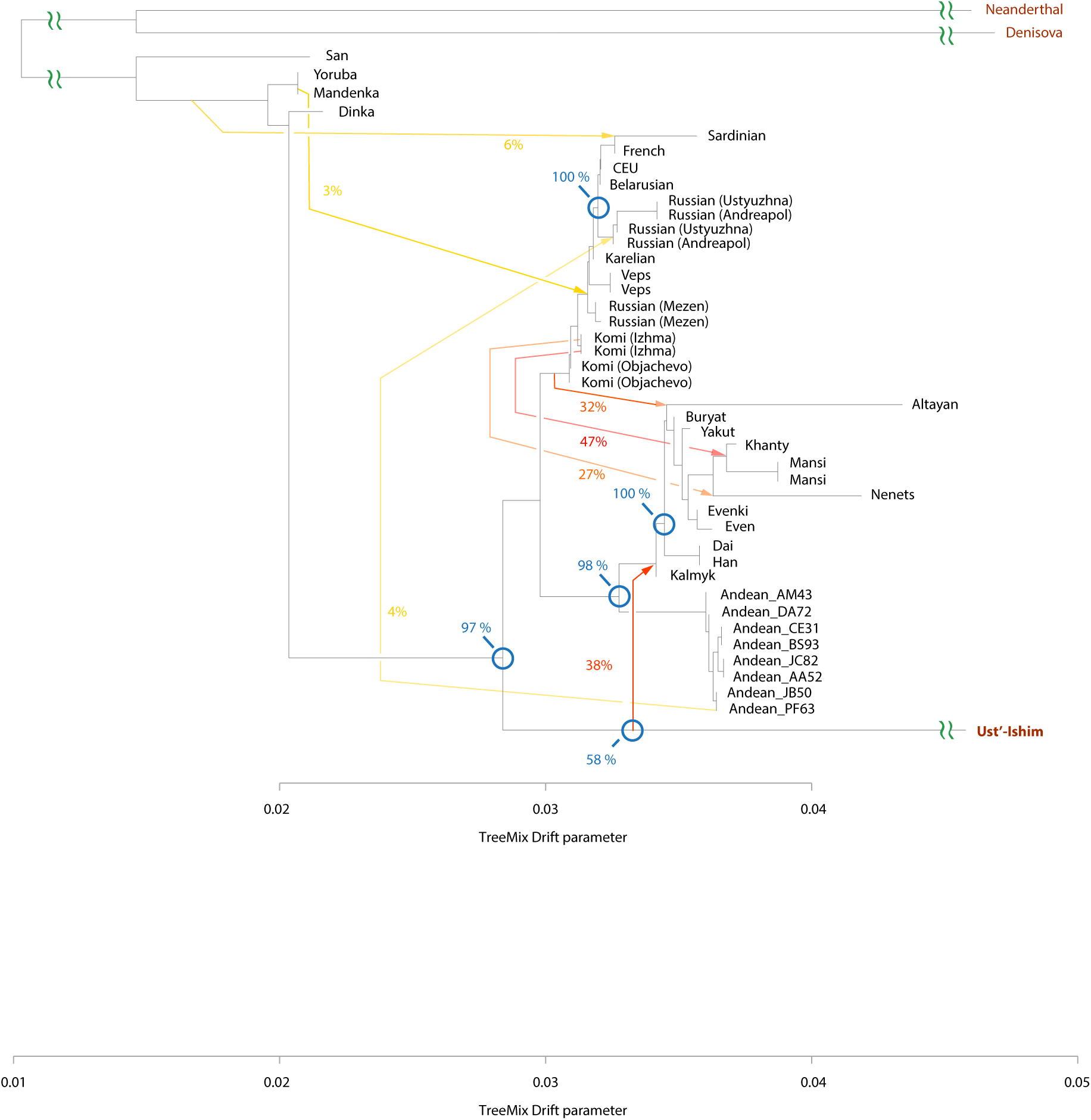
Genetic relationships of Ust’-Ishim and modern-day humans. TreeMix model with 7 admixture events incorporates ancient Siberian Ust’-Ishim. Blue circles indicate branching events that were investigated using 100 bootstraps and percentages of trees supporting the branch point are shown. Residuals are shown in Supp. Fig. 14.

Our data predicts that 41% (95% CI: 36-45%) of the ancestry of Andean Highlanders, the Native Americans, is attributable to the ANE lineage represented by ancient Siberians MA-1 and AG-2 (Fig. 7; Supp. Fig. 16). This agrees with previous reports that Paleo-americans trace approximately 42% of their autosomal genome to an ancient Eurasian lineage related to MA-1^5^. Surprisingly, our tree model also grouped Mansi and Nenets people together with the ancient individuals MA-1 and AG-2, demonstrating the genetic link between modern Western Siberians and ancient North Eurasians. We tested this prediction using the D-statistic, by evaluating whether a modern individual shared significantly more derived alleles with MA-1 or AG-2 than with the East Asian Han individual (Fig. 5d, Supp. Fig. 17a). Indeed, among non-Europeans Mansi, Khanty, Nenets and Native Americans all have very strong ANE ancestry, demonstrated by their significant genetic affinities with MA-1 and AG-2. Furthermore, the TreeMix model (Fig. 7) inferred that 43% of Mansi ancestry was derived from the admixture with a population related to Eastern Siberians Evenki and Even, while 57% is attributable to the ANE ancestry. The European-related ancestry signals within Mansi genomes are unlikely to be explained by a recent admixture of Mansi with Komi (Supp. Fig. 18), suggesting that Mansi are descendants of Siberian ANE people as well as Eastern Siberians with similar proportions of both ancestries.

**Figure 7.**
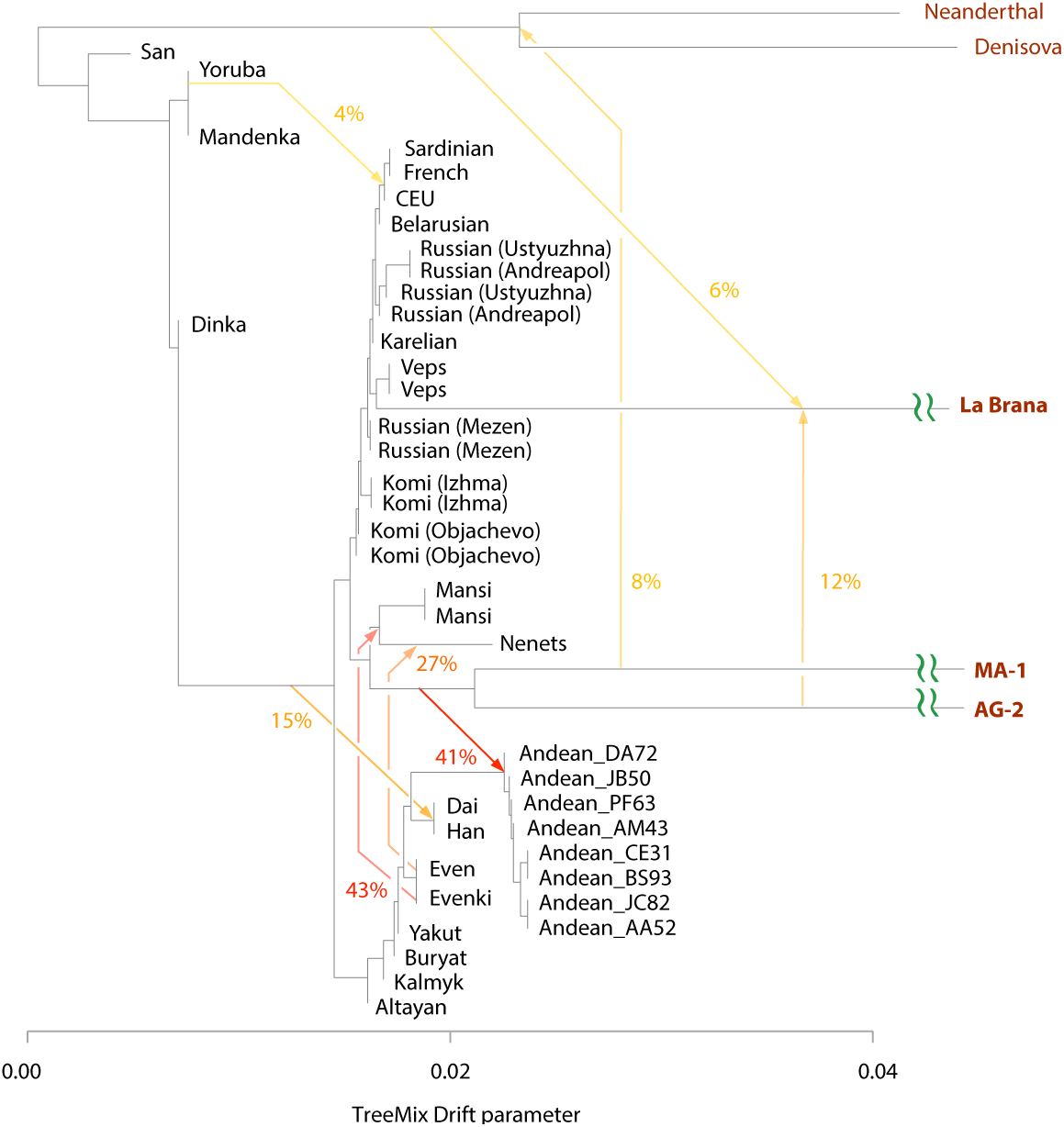
Genetic relationships with ancient North Eurasians MA-1 and AG-2. Autosomal TreeMix admixture graph that includes ancient 7,000-year old La Brana (Spain) and low-coverage genomes of ancient Siberians MA-1 and AG-2. Residual plot for this graph is shown in Supp. Fig. 16.

The tree model with the 4,000-year-old genome of Saqqaq from Greenland^34^ showed that Saqqaq was related to both East Asians and Eastern Siberians, and 12% of its ancestry was shared with the ancestral population of Even (Supp. Fig. 19-20). The D-statistic (Supp. Fig. 7e, 8e) also demonstrated the strong genetic affinity between Saqqaq and the Even population.

We further analyzed SNP data from the Holocene Eastern European hunter-gatherers^44^ from the Samara and Karelia regions in modern Russia and Motala region in Sweden (8-6.6 kya) as well as more recent Bronze Age Yamnaya samples (5.3-4.7 kya). D-statistics (Supp. Fig. 21) demonstrated strong admixtures between Mansi and nearly all hunter-gatherers from Eastern Europe, particularly for Samara and Karelia samples. Slightly weaker, but significant affinities were present between Eastern Siberian Even and Eastern European hunter-gatherers. This was consistent with Mansi carrying Even-related admixture, which also permeated into ancient European populations. Yamnaya samples also showed statistically significant admixtures with Mansi and Even, but weaker compared to Samara HG, which indicates further dilution of eastern hunter-gatherer ancestry component of Yamnaya culture samples. Since Mansi-related admixtures are detectable within an ancient individual, who lived 8-6.6 kya, the ANE-related ancestry among Eastern European hunter-gatherers could be attributed to gene flows between population ancestral to Mansi and Eastern European hunter-gatherers that occurred before 6,600 years ago. The unexpected genetic link between Mansi and ancient Hungarians may explain similarities between Mansi and Hungarian languages, although further analyses would be required to definitively establish this connection.

Siberians also shared part of their ancestry with Pitted Ware Culture (PWC) 5,000-year-old hunter-gatherers from Sweden Ire8 and Ajv52^33^. Ajv52 and particularly Ire8 had strong admixture signals with Western Siberians Mansi (Fig. 5e, Supp. Fig. 17b), suggesting that like the closely related Yamnaya culture, they had strong ANE ancestries likely due to admixtures with Mansi-related population.

## DISCUSSION

In this work we examined genetic history of several Siberian and Eastern European populations. We have summarized our findings in a geographical dispersion model (Fig. 8), which details how populations and genetic variation likely spread across Northern Eurasia.

**Figure 8.**
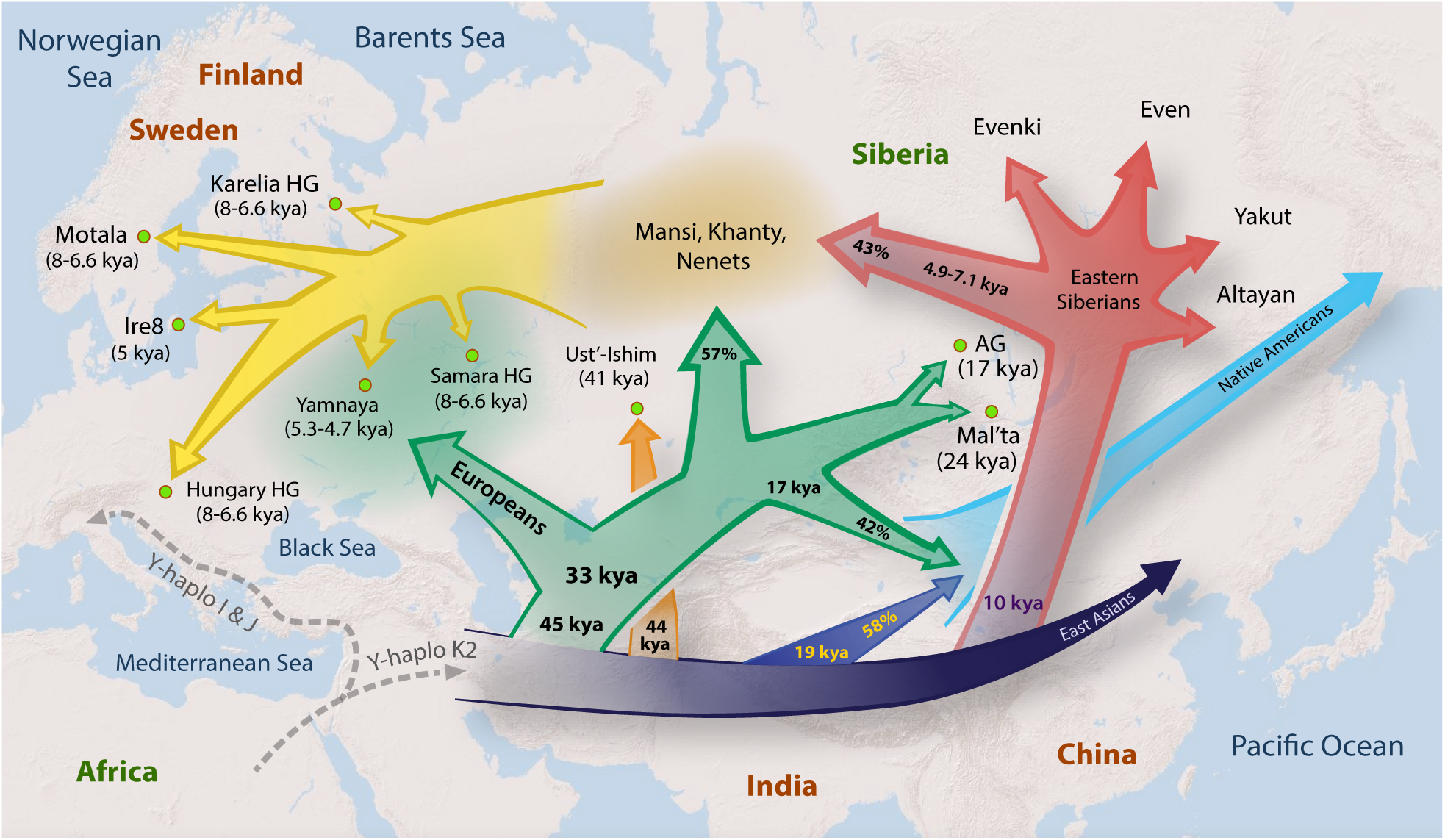
Geographical dispersion model. The approximate model of gene flows reflecting divergence of primary clades in Europe, East Asia and Siberia with approximate divergence times based on Chromosome Y, MSMC, TreeMix and D-statistics analyses. Percentages represent proportions of ancestry contributed to each lineage from other lineages. The common ancestors of Mansi, Khanty and Nenets had derived 57% of their ancestry from ANE population, related to MA-1, AG-2 and Native Americans and 43% from the admixture with a population related to Even and Evenki. Native Americans trace 42% of their ancestry to ANE and 58% to a common ancestor Eastern Siberians and East Asians. 45 kya represents the split between Y-DNA clades QR and NO. 33 kya represents the split betweenY-DNA Q and R clades and MtDNA N2 and W clades. 17 kya represents Mansi-Andean separation time. 44 kya represents the split of Ust’-Ishim’s haplogroup from NO clade. 19 kya represents Han-Andean separation time. 10 kya represents Han-Eastern Siberian separation time. 7.1 kya represents N-clade expansion. 4.9–7.1 kya represents Evenki-Mansi MSMC separation time (6.9 kya) and expansion of N haplogroup (4.9–7.1 kya). In this Figure Europeans represent a collective term for the following populations from this study: Komi, Veps, Karelians, Russians, and Belarussians.

Our analyses suggest that modern-day Eurasians diverged from ancestral African populations approximately 70,000 years ago, based on individual estimates of 69,000 (Y-DNA), 55,000 (autosomal) and 85,000 years (mtDNA). Divergence time estimates of major Y-DNA haplogroups indicate that some of the earliest Eurasian lineages such as C and H diverged as early as 69,000 and 50,000 years ago, followed by the formation of clades common to Europeans (R) and East Asians (O) 45,000 year ago. However, our autosomal MSMC analysis inferred that the largest separation time between the two modern-day Eurasians was only 28,000 (French and Han), suggesting that apparent genetic differences between modern-day Eurasians are significantly lower than would be expected from isolated populations.

Our results suggest that Eastern Siberians form a separate lineage, related to East Asian populations of Han and Dai with separation time of 8.8-11.2 kya. The existence of Eastern Siberian lineage, and its affinity to East Asian populations was well supported by multiple TreeMix models (Fig. 2, 6, 7, Supp. Fig. 3) as well as 100% of bootstrap replicates (Fig. 6).

The TreeMix model (Fig. 6) suggests that Eastern Siberian and East Asian populations shared a surprisingly large amount of ancestry (38%) attributable to ancient Siberian Ust’-Ishim, pointing to the very ancient origins of indigenous Siberians. However, the degree of shared ancestry with Ust’-Ishim should be interpreted with caution, given the multiple assumptions of the TreeMix models, such as instantaneous gene flows underlying admixture events and small amounts of genetic drift between populations. The shared ancestry of Eastern Asians, Siberians and Ust’-Ishim is also supported by the D-statistic (Fig. 5c, Supplementary Fig. 13c, Supp. Fig. 15), which showed that modern Eastern Asians and Eastern Siberians share more alleles with Ust’-Ishim compared to modern Europeans. Another line of evidence pointing to affinity of East Asians and Eastern Siberians with Ust’-Ishim was given by the Y-chromosome haplogroup analysis (Supp. Fig. 12). We showed that Ust’-Ishim was a member of the NO clade, predating the split of between Eastern/Southern Asian major Y-DNA haplogroups O and N, which our data suggests is of Eastern Siberian origin. This indicates that the split between N and O clades likely occurred in Siberia or Asia, rather than in Europe. Therefore, based on the new genome-wide data, our work provides a more accurate interpretation of patterns observed in the original Ust’-Ishim study^11^.

Our analysis indicates that Ust’-Ishim lived approximately 41,000 years ago. However, the estimated age of Ust’-Ishim depends on the mutation rate of Y-chromosome, that is not yet definitively established. If this parameter is changed, the split times of all haplogroups would scale up or down accordingly by the same percentage. Therefore our results indicate a potential discrepancy of approximately 10% between ages inferred using Y-chromosome mutation rate and ages of fossil bones inferred using chemical methods. Therefore, further systematic studies would be necessary to reconcile the estimates produced by the two methods.

The large-scale TreeMix analysis, which was free of ascertainment biases, revealed several previously unknown admixture events. We observed particularly robust admixture (5%) between unknown African population and East Asians Han, Dai, Sherpa and Malaysians (Fig. 2, Fig. 7, Supp. Fig. 19), consistent with survival of multiple human lineages in East Asia^45^, which contributed to genetic makeup of modern-day Asian populations. These results were also supported by the D-statistic analyses (Supp. Fig. 22), which showed that Dinka and Mbuti likely harbor signals of admixture with East Asians after they diverged from European populations represented by French, Belarusian or CEU (Z=0.8-2.2). However, these results should be interpreted with caution since the tests did not reach statistical significance of Z>=3. More in-depth analyses would be required to identify the specific African populations that could have contributed specifically to the East Asian ancestry.

A particularly surprising finding was the genetic link between 24,000-year-old Siberian Mal’ta boy, Native Americans and Western Siberians Mansi, Khanty and Nenets. We estimate that 57% of Mansi and Khanty ancestry are related to ANE (MA-1, AG-2). The D-statistics (Fig. 5d) agrees with this observation and shows that Western Siberians and Native Americans share a similar amount of ANE-related (MA-1, AG-2) ancestry (Fig. S17). The remaining 43% of Western Siberian ancestry is shared with Eastern Siberians, suggesting admixtures from Eastern Siberian populations most related to Even and Evenki into the common ancestors of Western Siberians (Fig. 2). The ANE-related ancestry is also significant among Northeastern European populations such as Mezen Russians, Komi, Karelians and Veps (Fig. 5d), suggesting their shared ancestry with Western Siberians. These northeastern European populations are also admixed with Eastern Siberians such as Yakut, Buryat and Even (Fig. 5a-b, Supp. Fig. 7f,j,k). Therefore some northeastern European populations are related to both ANE and Eastern Siberians. These observations are consistent with previous findings^4^ that ancestry of Eastern Europeans such as Mordovians, Finns, Russians, Saami and Chuvash cannot be explained by a mixture of three early European groups: the ancient northern Eurasians (ANE, represented by MA-1), the West European hunter-gatherers (WHG, represented by Loschbour), and the early European farmers (EEF, represented by Stuttgart). These Eastern European groups are more related to East Asians^4^, which agrees with our observations of strong Eastern Siberian admixtures contributing to the ancestry of Northeastern Europeans.

Our findings also suggest that Western Siberians Mansi and Khanty and ancient eastern European populations share a significant amount of ANE-related ancestry. Indeed, Yamnaya culture people (5.3-4.7 kya) had stronger genetic affinity to ANE than Mansi or Finns to ANE (Supp. Fig. 21a). Yamnaya people shared more alleles with Mansi than Western Europeans French and Sardinian (Supp. Fig. 21b), supporting shared ANE ancestry among Mansi and Yamnaya people. And importantly, Yamnaya people have greater affinity to Mansi than to Eastern Siberian population of Even (Supp. Fig. 21c), favoring Western Siberians over Eastern Siberians as specifically contributing to the Yamnaya ancestry. The Pitted Ware culture Ire8 and Ajv52 samples from Sweden, who are genetically related to the Yamnaya people^44^, also harbor similarly high admixtures with Mansi (Fig. 5e, Supp. Fig. 24) and weak affinity with Eastern Siberians.

The Western Siberian admixture into the Eastern Europeans likely began before the Yamnaya culture period (5.3-4.7 kya), since the admixtures with Mansi are also very strong among hunter gatherers from Northeastern Europe from 6.6-8 kya (Karelia HG, Samara HG and to lesser degree Motala HG and Hungary Gamba HG; Supp. Fig. 21f-q) that predated the Yamnaya people. Therefore Western Siberian admixtures into northeastern Europe likely began prior to 6,600 years ago, coinciding with the expansion of Y-DNA haplogroup N1c1 among Siberians and northeastern Europeans (7,100-4,900 years ago). Since haplogroup N likely originates in Asia or Siberia, its presence among eastern Europeans likely reflects ancient gene flows from Siberia into Eastern Europe.

The rapid dispersal of haplogroup N1c1 7,100-4,900 years ago among Siberians and northeastern Europeans is surprising, given the significant geographical distance between Eastern Siberian and Eastern Europe. Such rapid movement of people may be explained by a technological breakthrough, such as invention of sleds or domestication of reindeer or horses, which enabled Siberians to rapidly move across vast expanses of Siberia and reach Eastern Europe, where they likely admixed with indigenous Europeans. Our work therefore establishes ancient genetic links of Siberian, European and Asian populations across 50,000 years.

## DATA AVAILABILITY

All sequencing data and SNP calls from the individuals that were sequenced as part of this study were deposited into the Sequence Read Archive (SRA) under the accession number PRJNA267856. Genotyping data was deposited to GEO (GSE70063).

## ACKNOWLEDGEMENTS

We would like to thank A. Sidow, J. Pickrell, C. Jeong, M. Rasmussen, P. Ralph, M. Ogneva and E. Lam for the insightful discussions and advice on the data analysis; V. Rabinovich, D. Gerasimova and R. Kuchin for their help with collecting Mansi and Khanty samples; M. Crawford, R. David, D. Van Den Berg, S. Tyndale, C. Nicolet, J. Herstein, J. Nguyen, P. Martinez, T. Michurina and G. Enikolopov for their help with DNA sequencing; Z. Albertyn and C. Hercus for their help with Novoalign; R. Ronen, V. Bafna, W. L. Ping, T. Y. Ying and T. K. Yong for facilitating exchange of sequencing data; S. Kim and J. Genovese for legal support; P. Thomas for insightful discussions.

J.N. would like to thank NIH (grant R01 HG007089). S.L. would like to thank grants from the Russian Basic Research Foundation (grant 13-04-00588); the Programs ‘Molecular and Cell Biology’ and ‘Fundamental Science for Medicine’ of the Russian Academy of Sciences; the Federal Support of Leading Scientific Schools (grant 2017.2014.4).

## REFERENCES

1 Genomes Project, C. et al. An integrated map of genetic variation from 1,092 human genomes. Nature 491, 56–65, doi:10.1038/nature11632 (2012).

2 International HapMap, C. et al. Integrating common and rare genetic variation in diverse human populations. Nature 467, 52–58, doi:10.1038/nature09298 (2010).

3 Der Sarkissian, C. et al. Ancient DNA reveals prehistoric gene-flow from siberia in the complex human population history of North East Europe. PLoS genetics 9, e1003296, doi:10.1371/journal.pgen.1003296 (2013).

4 Lazaridis, I. et al. Ancient human genomes suggest three ancestral populations for present-day Europeans. Nature 513, 409–413, doi:10.1038/nature13673 (2014).

5 Raghavan, M. et al. Upper Palaeolithic Siberian genome reveals dual ancestry of Native Americans. Nature 505, 87–91, doi:10.1038/nature12736 (2014).

6 Waters, M. R., Forman, S. L. & Pierson, J. M. Diring Yuriakh: A lower paleolithic site in central Siberia. Science 275, 1281–1284 (1997).

7 Hublin, J. J. Out of Africa: modern human origins special feature: the origin of Neandertals. Proceedings of the National Academy of Sciences of the United States of America 106, 16022–16027, doi:10.1073/pnas.0904119106 (2009).

8 Hublin, J. J. The earliest modern human colonization of Europe. Proceedings of the National Academy of Sciences of the United States of America 109, 13471–13472, doi:10.1073/pnas.1211082109 (2012).

9 Prufer, K. et al. The complete genome sequence of a Neanderthal from the Altai Mountains. Nature 505, 43–49, doi:10.1038/nature12886 (2014).

10 Meyer, M. et al. A high-coverage genome sequence from an archaic Denisovan individual. Science 338, 222–226, doi:10.1126/science.1224344 (2012).

11 Fu, Q. et al. Genome sequence of a 45,000-year-old modern human from western Siberia. Nature 514, 445–449, doi:10.1038/nature13810 (2014).

12 McKenna, A. et al. The Genome Analysis Toolkit: a MapReduce framework for analyzing next-generation DNA sequencing data. Genome research 20, 1297–1303, doi:10.1101/gr.107524.110 (2010).

13 Purcell, S. et al. PLINK: a tool set for whole-genome association and population-based linkage analyses. American journal of human genetics 81, 559–575, doi:10.1086/519795 (2007).

14 Patterson, N., Price, A. L. & Reich, D. Population structure and eigenanalysis. PLoS genetics 2, e190, doi:10.1371/journal.pgen.0020190 (2006).

15 Price, A. L. et al. Principal components analysis corrects for stratification in genome-wide association studies. Nature genetics 38, 904–909, doi:10.1038/ng1847 (2006).

16 Fedorova, S. A. et al. Autosomal and uniparental portraits of the native populations of Sakha (Yakutia): implications for the peopling of Northeast Eurasia. BMC evolutionary biology 13, 127, doi:10.1186/1471-2148-13-127 (2013).

17 Zhou, D. et al. Whole-genome sequencing uncovers the genetic basis of chronic mountain sickness in Andean highlanders. American journal of human genetics 93, 452–462, doi:10.1016/j.ajhg.2013.07.011 (2013).

18 Li, J. Z. et al. Worldwide human relationships inferred from genome-wide patterns of variation. Science 319, 1100–1104, doi:10.1126/science.1153717 (2008).

19 Khrunin, A. V. et al. A genome-wide analysis of populations from European Russia reveals a new pole of genetic diversity in northern Europe. PloS one 8, e58552, doi:10.1371/journal.pone.0058552 (2013).

20 Yunusbayev, B. et al. The Caucasus as an asymmetric semipermeable barrier to ancient human migrations. Molecular biology and evolution 29, 359–365, doi:10.1093/molbev/msr221 (2012).

21 Wong, L. P. et al. Insights into the genetic structure and diversity of 38 South Asian Indians from deep whole-genome sequencing. PLoS genetics 10, e1004377, doi:10.1371/journal.pgen.1004377 (2014).

22 Wong, L. P. et al. Deep whole-genome sequencing of 100 southeast Asian Malays. American journal of human genetics 92, 52–66, doi:10.1016/j.ajhg.2012.12.005 (2013).

23 Jeong, C. et al. Admixture facilitates genetic adaptations to high altitude in Tibet. Nature communications 5, 3281, doi:10.1038/ncomms4281 (2014).

24 Pickrell, J. K. & Pritchard, J. K. Inference of population splits and mixtures from genome-wide allele frequency data. PLoS genetics 8, e1002967, doi:10.1371/journal.pgen.1002967 (2012).

25 Erdeniev, U. E. Kalmyks. (Moscow: Nauka., 1985).

26 Lipson, M. et al. Reconstructing Austronesian population history in Island Southeast Asia. Nature communications 5, 4689, doi:10.1038/ncomms5689 (2014).

27 Durand, E. Y., Patterson, N., Reich, D. & Slatkin, M. Testing for ancient admixture between closely related populations. Molecular biology and evolution 28, 2239–2252, doi:10.1093/molbev/msr048 (2011).

28 Patterson, N. et al. Ancient admixture in human history. Genetics 192, 1065–1093, doi:10.1534/genetics.112.145037 (2012).

29 Schiffels, S. & Durbin, R. Inferring human population size and separation history from multiple genome sequences. Nature genetics 46, 919–925, doi:10.1038/ng.3015 (2014).

30 Poznik, G. D. et al. Sequencing Y chromosomes resolves discrepancy in time to common ancestor of males versus females. Science 341, 562–565, doi:10.1126/science.1237619 (2013).

31 Rasmussen, M. et al. The genome of a Late Pleistocene human from a Clovis burial site in western Montana. Nature 506, 225–229, doi:10.1038/nature13025 (2014).

32 Olalde, I. et al. Derived immune and ancestral pigmentation alleles in a 7,000-year-old Mesolithic European. Nature 507, 225–228, doi:10.1038/nature12960 (2014).

33 Skoglund, P. et al. Origins and genetic legacy of Neolithic farmers and hunter-gatherers in Europe. Science 336, 466–469, doi:10.1126/science.1216304 (2012).

34 Rasmussen, M. et al. Ancient human genome sequence of an extinct Palaeo-Eskimo. Nature 463, 757–762, doi:10.1038/nature08835 (2010).

35 Rasmussen, M. et al. An Aboriginal Australian genome reveals separate human dispersals into Asia. Science 334, 94–98, doi:10.1126/science.1211177 (2011).

36 Goebel, T., Waters, M. R. & O’Rourke, D. H. The late Pleistocene dispersal of modern humans in the Americas. Science 319, 1497–1502, doi:10.1126/science.1153569 (2008).

37 Underhill, P. A. et al. The phylogenetic and geographic structure of Y-chromosome haplogroup R1a. European journal of human genetics: EJHG 23, 124–131, doi:10.1038/ejhg.2014.50 (2015).

38 Khar’kov, V. N. et al. [The origin of Yakuts: analysis of Y-chromosome haplotypes]. Molekuliarnaia biologiia 42, 226–237 (2008).

39 Tambets, K. et al. The western and eastern roots of the Saami--the story of genetic “outliers” told by mitochondrial DNA and Y chromosomes. American journal of human genetics 74, 661–682, doi:10.1086/383203 (2004).

40 Lappalainen, T. et al. Migration waves to the Baltic Sea region. Annals of human genetics 72, 337–348, doi:10.1111/j.1469-1809.2007.00429.x (2008).

41 Malyarchuk, B. et al. Differentiation of mitochondrial DNA and Y chromosomes in Russian populations. Human biology 76, 877–900 (2004).

42 Mirabal, S. et al. Y-chromosome distribution within the geo-linguistic landscape of northwestern Russia. European journal of human genetics: EJHG 17, 1260–1273, doi:10.1038/ejhg.2009.6 (2009).

43 Lappalainen, T. et al. Regional differences among the Finns: a Y-chromosomal perspective. Gene 376, 207–215, doi:10.1016/j.gene.2006.03.004 (2006).

44 Haak, W. et al. Massive migration from the steppe was a source for Indo-European languages in Europe. Nature 522, 207–211, doi:10.1038/nature14317 (2015).

45 Chang, C. H. et al. The first archaic Homo from Taiwan. Nature communications 6, 6037, doi:10.1038/ncomms7037 (2015).

